# Ecdysteroids as natural doping substances in the blood of insectivorous bats

**DOI:** 10.1101/2021.05.12.443761

**Authors:** Sándor Hornok, Róbert Berkecz, Endre Sós, Attila D. Sándor, Tímea Körmöczi, Norbert Solymosi, Jenő Kontschán, Attila Hunyadi

## Abstract

Bats have deserved much scientific attention due to their biological-ecological properties and increasingly recognized epidemiological significance. Ecdysteroids are insect-molting hormones that (under experimental conditions) have stimulating and anabolic effects in mammals, including humans. Therefore, these biologically active compounds are currently under consideration by the World Anti-Doping Agency (WADA) to become doping-controlled substances. Previously we demonstrated that low to high concentrations of ecdysteroids appear in the blood of insectivorous passerine birds. Since passerine birds and echolocating bats share several adaptive mechanisms in connection with flying, and insectivorism is also among their common traits, we hypothesized that ecdysteroids might also be present in the blood of insectivorous bats. To test this hypothesis, blood samples of eight insectivorous bat species were collected and analyzed for the presence of ecdysteroids with highly sensitive targeted ultra-high-performance liquid chromatography coupled to high-resolution quadrupole-orbitrap mass spectrometry method (UHPLC-HRMS). The results supported our hypothesis, because nine ecdysteroids were detected in bat blood. The spectrum of these ecdysteroids was similar in those bat species which have their most preferred food items from the same insect order, supporting insects as the most likely source of these hormones. It was also shown that the spectrum of blood-borne ecdysteroids was broader in the autumn than in the summer, and higher concentrations of 20-hydroxyecdysone were measured in samples of large size bat species in comparison with small size ones. Based on the known physiologic effects of ecdysteroids, we postulate that these results might have implications on the metabolic rate and parasite burdens of insectivorous bats.

---

Bats (Mammalia: Chiroptera) have attracted enormous scientific attention on account of their biological-ecological properties and epidemiological significance. Importantly, they are the only mammals that actively fly. This adaptation to flying entailed unique anatomical and physiological mechanisms, including a high metabolic rate^1^. Insects, on which the great majority of bat species feed, contain molting hormones, so-called ecdysteroids. Recently it has been demonstrated that low to high concentrations of ecdysteroids appear in the blood of insectivorous passerine birds and these might possibly contribute to their “higher-than-avian-average” metabolic rate and physical performance^2,3^. Because passerine birds and insectivorous bats share several adaptive mechanisms in connection with flying, and dietary guild is also among their common traits, we hypothesized that ecdysteroids may also be present in the blood of insectivorous bats. The aim of this study was to investigate this possibility.

In the blood of eight insectivorous bat species, sampled over a one-year period, nine ecdysteroids were detected (Figure 1; Supplementary Table 1). Ecdysteroids were the most diverse in *Eptesicus serotinus*, followed by *Nyctalus noctula* and *Pipistrellus nathusii* (Figure 1). Considering the whole study period, the spectra of ecdysteroids were identical or overlapping between bat species which have their most preferred food items from the same insect order (Technical Appendix). In particular, all compounds detected in *Hypsugo savii* were also present in *Miniopterus schreibersii* (preferentially feeding on Lepidopterans), all compounds shown to be present in *Myotis myotis* were also demonstrated in *E. serotinus* (preferentially feeding on beetles), and *My. nattereri* shared all its compounds with *N. noctula* and *P. nathusii* (Figure 1). In case of the latter two species the spectrum of ecdysteroids was the same, in accordance with their most preferred food items, Dipterans, but despite of the highly significant difference between their mean body weight (Supplementary Table 2: 28.1±3.63 g, n=192 *vs* 6.76±1.48 g, n=44, respectively) (t=62.015, df=170.38, P<0.0001). The mean number of compounds per sampling occasion was low in January, April and compared to the summer (median=4, IQR=1.25, n=20 occasions) it was significantly (t=2.34, df=26, P=0.028) higher during the autumn (median=6, IQR=2.5, n=7 occasions) (Supplementary Table 1).

**Figure 1.**
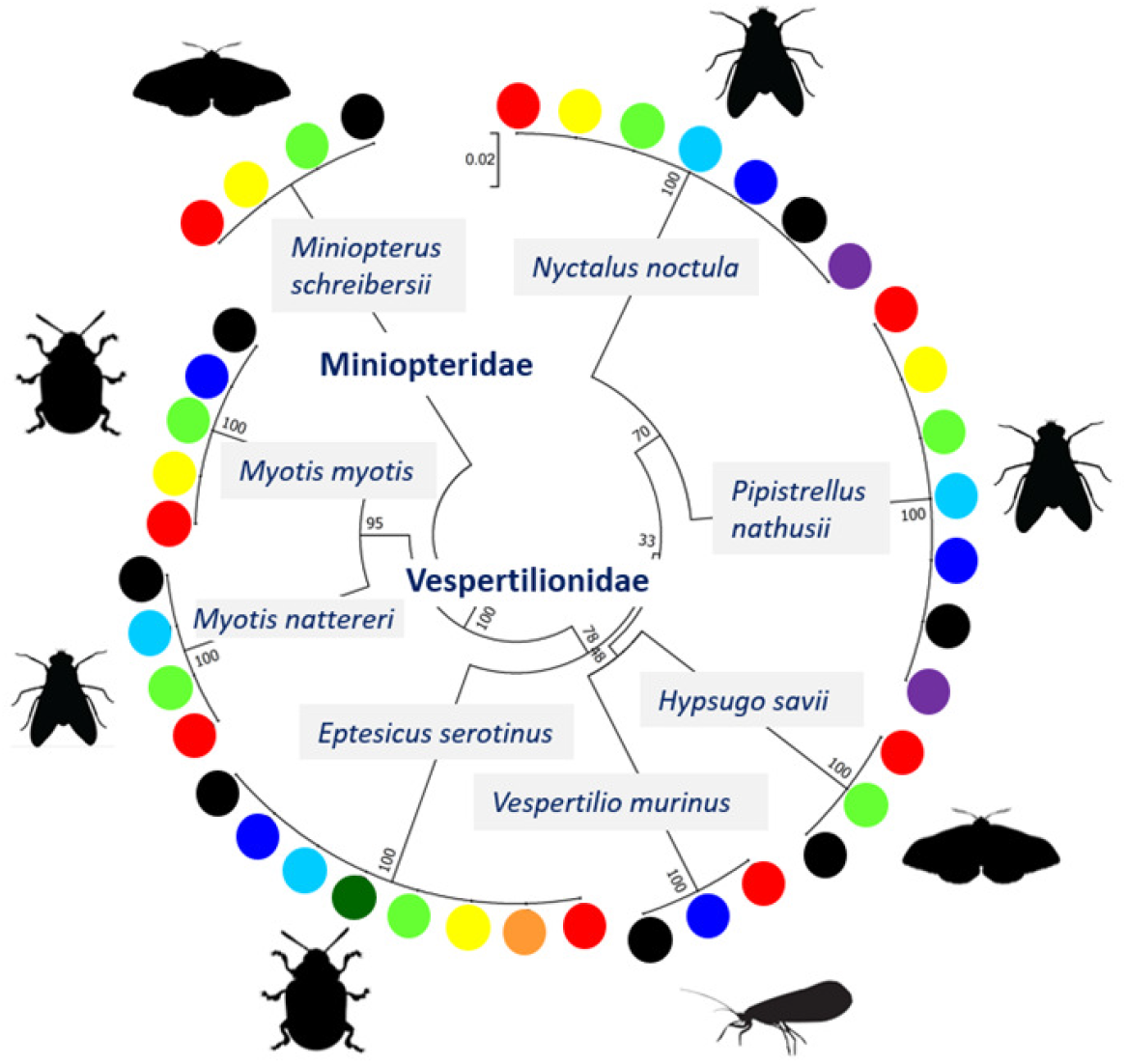
Diversity of ecdysteroids in bat species according to their most preferred food items. The topology is based on a cytochrome *b* phylogenetic tree. The color codes of ecdysteroids are: red – 20-hydroxyecdysone, orange – polypodine B, yellow – poststerone, light green – 14-deoxy-20-hydroxyecdysone, dark green – ecdysone, light blue – ajugasterone C, dark blue – calonysterone, black – dacryhainansterone, purple – shidasterone. The pictograms of food items stand for beetles (order Coleoptera), moths (order Lepidoptera), flies (mosquitoes also included: order Diptera) and caddisflies (order Trichoptera).

The most prevalent ecdysteroid (detected in all evaluated bat individuals) was 20-hydroxyecdysone (20E). Among the three large size bat species (*N. noctula, E. serotinus*, and *My. myotis*), the mean 20E concentration was significantly higher in *E. serotinus* (232.82±287.23 pmol/ml, n=8) compared to *N. noctula* (38.49±27.92 pmol/ml, n=11) (W=77, P=0.012). In addition, the mean concentration of 20E was significantly higher in bat species predominantly feeding on beetles (order Coleoptera) (204.96±260.09, n=10) than in those typically feeding on flies (order Diptera) (49.74±59.58, n=15) (W=129, P=0.009) (Figure 1).

Thus, in the blood of all eight insectivorous bat species studied here the presence of a broad range of naturally acquired ecdysteroids was demonstrated, to the best of our knowledge for the first time in a worldwide context. Hitherto the general view persisted that experimentally administered ecdysteroids are rapidly cleared from the blood of mammals, e.g., mice^4^. However, the continuous uptake of insects can apparently counterbalance this rapid clearance in bats.

Regarding the physiologic importance of these findings, various ecdysteroids were shown to have significant anabolic effect in mammals, sometimes exceeding that of the well-known doping agents. For instance, 20E enhances physical performance, acting as a “natural dope”,^5^ and it is currently under consideration by the World Anti-Doping Agency to become a doping-controlled substance. Based on such data, it can be hypothesized that bats may take advantage of these biologically active compounds in their blood, particularly in association with their high metabolic rate necessitated by flying.

The predominant source of ecdysteroids detected in bats should have been insects, considering that all eight bat species investigated here are insectivorous. While none of the evaluated bat species are known to feed on fruits, they may still have access to plant-derived ecdysteroids via their insect prey items which feed on plants. This should explain the surprising occurrence of phytoecdysteroids in bat blood as shown here (i.e., those which are not known to be naturally produced by insects, such as polypodine B, ajugasterone C, calonysterone, dacryhainansterone and shidasterone).

Based on our results, identical spectra of detected ecdysteroids in *N. noctula* and *P. nathusii* was in accordance with their common most preferred food items (Figure 1), but not with their significantly different body weight. Assuming that the spectrum of blood-borne ecdysteroids is interrelated with the taxonomic diversity of insect food items, these results confirm previous observations, according to which dietary diversity was not related to the body mass of bats^6^. At the same time, variations in the spectra of ecdysteroids according to sampling months might reflect changes in the daily or seasonal availability/uptake of insects from various taxonomic orders.

Based on the dataset of this study, within the category of large size bats, higher 20E concentrations were measured in *E. serotinus* than in *N. noctula. Eptesicus serotinus* (unlike *N. noctula*) belongs to the category of “ground-gleaning” bats^7^, i.e., obtains a significant portion of its food items from surface objects such as the ground and its plant covering. From such surfaces *E. serotinus* predominantly feeds on beetles, which might significantly contribute to its high 20E blood concentrations. It was observed that in case of beetles the adult diapause is induced by the commencing short-day photoperiod in the autumn, concomitantly with an ecdysteroid peak value nearly as high as those found during metamorphosis^8^. In a broader context, this may also explain why the blood of bat species predominantly feeding on beetles contained significantly higher concentrations of 20E compared to those species which typically feed on flies.

The oral uptake of excess amounts of ecdysteroids by blood-sucking ectoparasites is known to affect them in several ways. Experimentally, these hormones have antifeedant effect and induce early salivary gland degeneration in ticks^9^ and were shown to naturally affect ticks taking a blood meal from birds with ecdysteroids in their blood^3^. In support of the possibility that a similar phenomenon might exist among insectivorous bats, these were shown to have significantly lower ectoparasite loads than sympatric fruit-eating bats^10^.

## Figure Legends

**Supplementary Figure 1.**
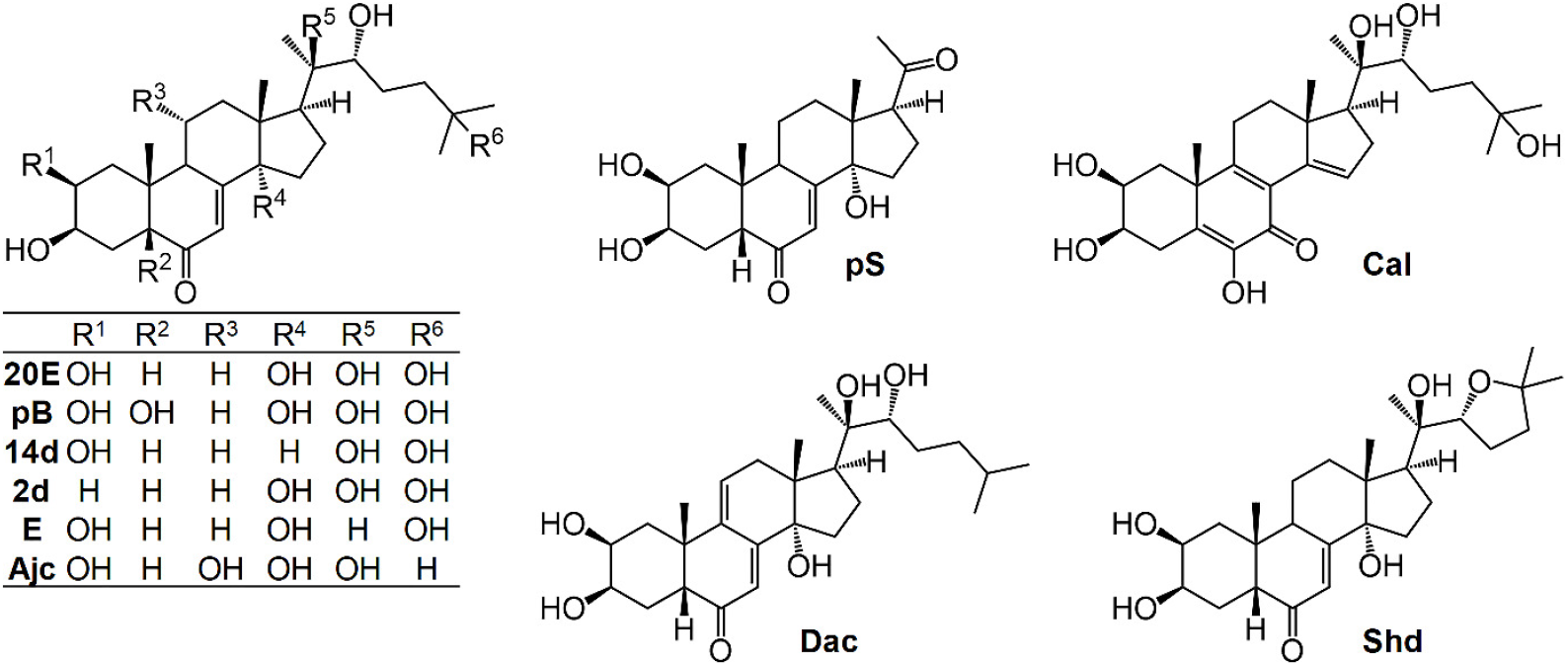
Chemical structures of the tested ecdysteroids. 20E – 20-hydroxyecdysone, pB – polypodine B, pS – poststerone, 14d – 14-deoxy-20-hydroxyecdysone, 2d – 2-deoxy-20-hydroxyecdysone, E – ecdysone, AjC – ajugasterone C, Cal – calonysterone, Dac – dacryhainansterone, Shd – shidasterone.

## Technical Appendix: Methods

### Origin of samples

In this study, blood samples of 36 bats were evaluated, representing eight insectivorous species from two families (Supplementary Table 1; Figure 1). The first group of blood samples (n=14) were collected from bats mist-netted at either a cave entrance (*My. myotis*, n=2) in Chavdartsi (Bulgaria), or near their roosts in buildings or mines at Canaraua Fetii, Romania (*N. noctula*: n=5; *P. nathusii*: n=2; *My. nattereri*: n=1 and *Mi. schreibersii*: n=4). Sampling dates included early September in 2017, the end of July and the beginning of August in 2018. All soft and hard ticks were removed from these bats and stored in 96% ethanol until identification according to standard keys^1^.

The second group of blood samples originated from 22 bats, which were transported to the Animal Rescue Centre of the Budapest Zoo, after found weak or in a moribund state in Budapest between mid-August, 2017 and early September, 2018 (i.e., in 2017: August, September, October, November, December; in 2018: January, April, August, September). In this category, all winter samples were obtained from the common noctule bat (*N. noctula*) which is known to have flying and foraging activity during winter months in Central Europe^2^, therefore its non-hibernating individuals could be included in our study from both December and January. All rescued bats were sampled as part of their veterinary care within one day of their arrival, therefore their blood ecdysteroids were regarded to originate from previous (thus natural) feeding. No ticks were found on these bats. Since no body mass data were available for bats in the second group, the mean weight of the species was calculated from a larger sample size encompassing the period 2015-2020 and including bats captured at various locations in Romania (Supplementary Table 2: A. D. Sándor, unpublished data).

### Chemicals and standards

All chemicals (water, methanol, acetonitrile, formic acid) for sample preparation and analysis with LC-MS grade were purchased from VWR (Radnor, PA, USA). Poststerone (20*E*)-oxime was used as internal standard (IS), and it was synthesized from poststerone as published before^3^.

The samples were tested for the presence of ten natural ecdysteroids, these compounds were obtained from our previous phytochemical work^4,5,6^. Abbreviations of these compounds’ trivial names are as follows: 20E – 20-hydroxyecdysone, pB – polypodine B, pS – poststerone, 14d – 14-deoxy-20-hydroxyecdysone, 2d – 2-deoxy-20-hydroxyecdysone, E – ecdysone, AjC – ajugasterone C, Cal – calonysterone, Dac – dacryhainansterone, Shd – shidasterone; chemical structures of these compounds are presented in Supplementary Figure 1.

### Sample preparation and calibration

The 150 µL of diluted blood sample (15 µL whole blood mixed with 135 µL saline solution) was spiked with 7 µL of IS solution (100 ng/mL in MeOH), and then 1 mL MeOH was added. After vortexing, the sample was shaken for 15 min at room temperature. Next, the sample was centrifuged at 4°C for 15 min at 21,000 g (Universal 320 R, Hettich, Tuttlingen, Germany) and 900 µL of the upper phase was dried under nitrogen at ambient temperature. The dried extract was dissolved in 65 µL of water/methanol (1/1, v/v) and centrifuged at 21,000 g at room temperature for 10 min, and the upper phase was collected for analysis.

External calibration was used for quantitative analysis of 10 ecdysteroids with the following concentrations of standards: 0, 0.033, 0.333, 1.667, 3.333, 16.667, 33.333, 333.333 and 3333.333 pmol/mL in water/methanol (1/1, v/v).

For calibration, standard solution mixtures were prepared with concentrations of 0, 0.033, 0.333, 1.667, 3.333, 16.667, 33.333 and 333.333 pmol/mL of each of pB, pS, 14d, E, AjC, Cal, Dac, 2d, and Shd in water/methanol (1/1, v/v). The concentration of 20E was set to 10 times higher than other standards and the concentration of IS was 28 pmol/mL in each calibration mixture.

### UHPLC-HRMS and HRMS/MS analysis

The UHPLC-HRMS and HRMS/MS measurements were performed on Waters Acquity I-Class UPLC™ (Waters, Milford, MA, USA) coupled to Thermo Scientific Q Exactive™ Plus Hybrid Quadrupole-Orbitrap™ (Thermo Fisher Scientific, Waltham, MA, USA) mass spectrometer. The UHPLC separation was carried out on an Accucore C30 column (150×2.1 mm, 2.6 µm) with an equivalent guard column (10×2.1 mm, 2.6 µm) from Thermo Fisher Scientific (Waltham, MA, USA). The mobile phase A consisted of 0.1% formic acid aqueous solution, and mobile phase B was composed of acetonitrile with 0.1% v/v formic acid. Total run time was 29 min and the following gradient program was used: 0 min 10% B held for 1.5 min; ramped to 35% B in 18.5 min; then ramped to 100% B in 0.1 min; held for 4.9 min; and finally, returned to initial conditions within 4 min. The flow rate was 0.4 mL/min during the analysis. Column temperature was maintained at 50 °C and the injection volume was 8 μL. Each sample was measured three times.

The mass spectrometer was operated in positive mode using single ion monitoring (SIM) for quantitative analysis of ecdysteroids and HRMS/MS parallel reaction monitoring (PRM) for their identification. The heated ESI source was used with the following conditions: capillary temperature 262.5 °C, S-Lens RF level 50, spray voltage 4.0 kV, sheath gas flow 50, spare gas flow 2.5, and auxiliary gas flow 12.50 in arbitrary units. For SIM mode the maximum injection time (IT) was 50 ms with a resolution of 35,000 (FWHM) and the automatic gain control (AGC) setting was defined as 1×10^6^ charges. In PRM mode with resolution of 17,500 (FWHM), the AGC setting was defined as 1×10^6^ charges, the maximum IT was set to 50 ms and 20 eV collision energy was used. The precursor ion window was set to 1 Da in both scan modes. The monitored m/z values of protonated compounds with related retention times are detailed in Supplementary Table 3.

The UHPLC system was controlled with MassLynx V4.1 SCN 901 (Waters, Milford, MA, USA). The control of mass spectrometer, data acquisition and processing were performed by Xcalibur 4.3 software (Thermo Fisher Scientific, Waltham, MA, USA).

In whole blood samples, the ecdysteroids were reliably confirmed by UHPLC-HRMS/MS analysis which were based on literature data and the fragmentation pattern of the reference standards^11^. The mass tolerance for the processing of UHPLC-HRMS spectra was set to 5 ppm. Three replicate quantitative analyses of samples were performed by external calibration using an internal standard and the linear calibration curves of ecdysteroids were based on analyte/IS peak area ratios against concentrations.

### Ethical approval

Permission for bat capture was provided by the Underground Heritage Commission (Romania) and the local protected area management authorities. Bat banding license numbers are 718/2017 (Bulgaria), 24/2017 and 111/2018 (Romania). Bats were handled according to the current law of animal welfare regulation (Law 206/2004), and the Research Bioethics Commission of USAMV CN approved the used methodology of bat handling and blood-collection protocol. Permission from the Institutional Animal Care and Use Committee (IACUC) was not necessary, because all wild-caught bats were released immediately in the field (none taken to participating institutes).

### Phylogenetic analysis

Circular phylogenetic tree was prepared to show the gross relationship of the eight bat species involved in this study (Figure 1) by including cytochrome *b* sequence data from GenBank^7^. This dataset was resampled 1,000 times to generate bootstrap values. Phylogenetic analyses were conducted with the Neighbor-Joining method, p-distance model, by using MEGA version 7.0.

### Statistical analyses

Statistically analyzed data included the concentrations of 20E (which was available in the case of all bats) compared between bat groups distinguished according to the following characters: bat species (*N. noctula vs E. serotinus*), body size characteristic of bat species (small species: mean body weight below 13 g, large species: mean body weight above 24 g; Supplementary Table 2), the predominant food item (from orders Coleoptera, Diptera, Trichoptera or Lepidoptera) characteristic of each bat species^8,9,10^. The mean concentrations of 20E were compared in male *vs* female bats, and between tick-infested and non-infested bats. In addition, the number of ecdysteroid compounds was analyzed according to astronomical seasons (summer sampling: July to early September; autumn sampling: October to early December). Where the assumption of normality was not satisfied, the Wilcoxon rank sum test^11^ was used, otherwise the *t*-test. Normality was verified by the Shapiro-Wilk test^12^. The P-values estimated by testing the groups on E20 were adjusted by the method of Benjamini and Hochberg^13^. The comparison of compound number between the summer and spring was performed using a negative binomial model^14^. All statistical analyzes were done in the R environment^15^. Differences were considered significant if P<0.05.

## Data availability

Important data are included in the manuscript and its references. Additional data are available in Supplementary Tables.

## Acknowledgments

This work was supported by the National Research, Development and Innovation Office, Hungary (NKFIH; K-119770, K-132794), the Economic Development and Innovation Operative Program GINOP-2.3.2-15-2016-00012, and the Ministry of Human Capacities, Hungary grant 20391-3/2018/FEKUSTRAT. Project no. TKP-32-1/PALY-2020 has been implemented with the support provided from the National Research, Development and Innovation Fund of Hungary, financed under the “Tématerületi Kiválósági Program 2020 (2020-4.1.1-TKP2020)” funding scheme. The authors are grateful to M. Vágvölgyi for preparing the ecdysteroid oxime used as internal standard, and to L. Barti, A. Corduneanu, I. Csősz and Á. Péter for invaluable help during fieldwork.

## Author contributions

S.H. initiated and designed the study, identified ticks, wrote most parts of the manuscript. R.B. and T.K. performed chemical analyses, E.S. and ADS provided expertise and important samples to the study, N.S. performed statistical analyses, J.K. performed phylogenetic analysis, A.H. supervised chemical analyses and wrote a significant part of the manuscript.

---

**Supplementary Table 1.** Ecdysteroid concentrations according to bat species, sex, sample group and sampling month.

| Supplementary Table 1. |  |  | Ecdysteroid concentrations according to bat species, sex, sample group and sampling month. |  |  |  |  |  |  |  |  |  |  |  |  |  |  |  |  |  |  |  |
| --- | --- | --- | --- | --- | --- | --- | --- | --- | --- | --- | --- | --- | --- | --- | --- | --- | --- | --- | --- | --- | --- | --- |
|  |  |  | Concentration (pmol/mL) |  |  |  |  |  |  |  |  |  |  |  |  |  |  |  |  |  |  |  |
| | | | 20E | | pB | | pS | | BKE11 | | $\alpha$ | | AjC | | Calo | | DeO-β | | Dakri | | Shida | |
| Species | Sex | Habitat (month) | pmol/mL | RSD% | pmol/mL | RSD% | pmol/mL | RSD% | pmol/mL | RSD% | pmol/mL | RSD% | pmol/mL | RSD% | pmol/mL | RSD% | pmol/mL | RSD% | pmol/mL | RSD% | pmol/mL | RSD% |
| PNAT | F | R (Aug) | 245,53 | 7,68 | ND |  | 273,41 | 2,6 | 130,71 | 7,43 | ND |  | 9,18 | 0,66 | 55,80 | 6,4 | ND |  | 9,53 | 8,1 | 5,32 | 16,3 |
| NNOC | M | R (no data) | 49,90 | 3,84 | ND |  | ND |  | 12,05 | 10,71 | ND |  | ND |  | 8,05 | 23,0 | ND |  | ND |  | ND |  |
| ESER | F | R (Aug) | 54,02 | 5,79 | ND |  | ND |  | 3,71 | 4,84 | ND |  | ND |  | 4,78 | 4,4 | ND |  | ND |  | ND |  |
| ESER | M | R (Oct) | 93,71 | 3,62 | ND |  | 49,27 | 9,0 | 7,43 | 4,66 | ND |  | 6,40 | 2,06 | 13,20 | 4,5 | ND |  | 4,49 | 4,3 | LOD |  |
| ESER | M | R (Oct) | 56,32 | 3,33 | 5,65 | 3,50 | 6,53 | 19,4 | 3,85 | 7,44 | 154,10 | 3,51 | 3,70 | 5,76 | 5,81 | 3,3 | ND |  | 4,90 | 2,0 | ND |  |
| no data | F | R (Oct) | 33,99 | 4,40 | ND |  | 24,48 | 5,2 | 11,34 | 3,31 | ND |  | 6,59 | 10,88 | 6,86 | 4,2 | ND |  | 17,84 | 8,4 | 5,78 | 7,5 |
| ESER | F | R (Nov) | 247,68 | 5,55 | 25,28 | 7,11 | 15,61 | 5,3 | 5,21 | 5,61 | LOD |  | LOD |  | 7,36 | 1,9 | ND |  | 5,37 | 10,3 | ND |  |
| NNOC | F | R (Dec) | 68,14 | 5,63 | ND |  | 14,24 | 8,8 | 9,60 | 6,25 | ND |  | 9,53 | 4,12 | 8,75 | 5,4 | ND |  | 37,85 | 4,0 | 4,38 | 14,8 |
| NNOC | F | R (Dec) | 54,23 | 6,81 | ND |  | 12,74 | 6,2 | 5,35 | 2,54 | ND |  | 4,75 | 9,10 | 8,98 | 8,2 | ND |  | 10,53 | 9,8 | ND |  |
| NNOC | F | R (Dec) | 5,96 | 13,09 | ND |  | 9,79 | 5,0 | LOD |  | ND |  | ND |  | ND |  | ND |  | ND |  | ND |  |
| MMYO | F | N (Sept) | 104,76 | 8,92 | ND |  | 12,51 | 28,8 | 7,12 | 18,01 | ND |  | LOD |  | LOD |  | ND |  | 5,98 | 22,1 | LOD |  |
| PNAT | M | N (Aug) | 34,29 | 5,00 | ND |  | 11,44 | 23,2 | 6,64 | 2,76 | ND |  | LOD |  | LOD |  | ND |  | 9,60 | 6,7 | ND |  |
| MSCH | F | N (Aug) | 45,21 | 2,64 | ND |  | 10,40 | 15,7 | 4,72 | 4,86 | ND |  | LOD |  | LOD |  | ND |  | 5,44 | 20,4 | LOD |  |
| MSCH | M | N (Aug) | 48,30 | 0,92 | ND |  | 19,76 | 4,8 | 5,61 | 3,46 | ND |  | LOD |  | LOD |  | ND |  | 14,86 | 3,5 | LOD |  |
| NNOC | M | N (July) | 49,94 | 5,84 | ND |  | 25,17 | 1,6 | 4,18 | 4,89 | ND |  | LOD |  | 72,80 | 3,8 | ND |  | 48,59 | 4,1 | LOD |  |
| NNOC | M | N (July) | 22,77 | 11,02 | ND |  | 11,35 | 4,4 | LOD |  | ND |  | ND |  | LOD |  | ND |  | LOD |  | ND |  |
| PNAT | M | N (Aug) | 8,31 | 1,27 | ND |  | LOD |  | ND |  | ND |  | ND |  | LOD |  | ND |  | ND |  | ND |  |
| VMUR | F | R (Apr) | 8,67 | 11,64 | ND |  | ND |  | ND |  | ND |  | ND |  | 43,38 | 2,6 | ND |  | LOD |  | ND |  |
| NNOC | M | N (July) | 16,66 | 16,86 | ND |  | ND |  | LOD |  | ND |  | ND |  | ND |  | ND |  | ND |  | ND |  |
| NNOC | M | R (Jan) | 3,01 | 11,19 | ND |  | ND |  | ND |  | ND |  | ND |  | ND |  | ND |  | ND |  | ND |  |
| ESER | M | R (no data) | 23,52 | 7,73 | ND |  | ND |  | LOD |  | ND |  | ND |  | ND |  | ND |  | 3,43 | 14,1 | ND |  |
| ESER | F | R (Aug) | 296,29 | 3,07 | 44,93 | 1,58 | ND |  | LOD |  | 250,01 | 2,47 | LOD |  | ND |  | ND |  | 6,80 | 10,9 | ND |  |
| HSav | M | R (Aug) | 48,08 | 5,52 | ND |  | LOD |  | 5,20 | 2,79 | ND |  | LOD |  | LOD |  | ND |  | 9,84 | 13,6 | ND |  |
| ESER | M | R (Sept) | 190,96 | 3,05 | ND |  | LOD |  | 10,65 | 18,31 | LOD |  | 21,28 | 9,12 | 12,60 | 18,1 | ND |  | 19,27 | 16,1 | LOD |  |
| no data | no data | R (no data) | 124,65 | 5,73 | LOD |  | LOD |  | LOD |  | ND |  | LOD |  | ND |  | ND |  | LOD |  | ND |  |
| no data | no data | R (no data) | 62,05 | 3,97 | ND |  | LOD |  | 6,88 | 9,97 | ND |  | 5,27 | 19,71 | 9,09 | 5,4 | ND |  | 14,86 | 16,6 | LOD |  |
| no data | no data | R (no data) | 13,54 | 4,21 | ND |  | ND |  | ND |  | ND |  | LOD |  | ND |  | ND |  | LOD |  | ND |  |
| ESER | no data | R (no data) | 900,08 | 0,16 | 12,78 | 3,50 | ND |  | 4,39 | 7,21 | 40,98 | 0,48 | ND |  | ND |  | ND |  | 5,27 | 33,8 | ND |  |
| NNOC | F | R (Nov) | 47,24 | 3,39 | ND |  | ND |  | LOD |  | ND |  | LOD |  | ND |  | ND |  | LOD |  | 7,07 | 19,4 |
| MMYO | F | N (Sept) | 82,22 | 2,95 | ND |  | LOD |  | 8,62 | 4,63 | ND |  | ND |  | 18,84 | 5,2 | ND |  | 5,51 | 9,1 | ND |  |
| MNAT | F | N (Aug) | 34,60 | 8,50 | ND |  | LOD |  | 6,72 | 17,87 | ND |  | 4,19 | 30,17 | ND |  | ND |  | 9,98 | 11,8 | ND |  |
| MSCH | M | N (Aug) | 8,34 | 30,64 | ND |  | LOD |  | 5,37 | 5,10 | ND |  | ND |  | ND |  | ND |  | 3,52 | 5,8 | ND |  |
| MSCH | M | N (Aug) | 8,90 | 32,97 | ND |  | LOD |  | ND |  | ND |  | ND |  | ND |  | ND |  | 4,07 | 3,9 | ND |  |
| NNOC | M | N (July) | 15,24 | 7,83 | ND |  | LOD |  | 6,46 | 4,08 | ND |  | ND |  | ND |  | ND |  | 3,88 | 16,0 | ND |  |
| NNOC | M | N (July) | 91,07 | 5,15 | ND |  | LOD |  | 14,70 | 6,64 | ND |  | 12,68 | 8,51 | ND |  | ND |  | 24,97 | 5,1 | ND |  |
| VMUR | M | R (Sept) | 12,06 | 3,71 | ND |  | ND |  | LOD |  | ND |  | LOD |  | ND |  | ND |  | 5,33 | 7,2 | ND |  |

**Supplementary Table 2.** Body weights (shown in grams) of bat species involved in this study, calculated from 2046 bats captured at various locations in Romania in the period 2015-2020.

|  | N | Minimum | Maximum | Mean | Std.<br>Deviation |
| --- | --- | --- | --- | --- | --- |
| *Nyctalus noctula* | 192 | 18,79 | 39,60 | 28,0968 | 3,62792 |
| *Pipistrellus nathusii* | 44 | 4,10 | 9,80 | 6,7632 | 1,48102 |
| *Hypsugo savii* | 6 | 8,60 | 11,00 | 9,8667 | ,98319 |
| *Vespertilio murinus* | 143 | 8,40 | 20,60 | 12,4116 | 2,29530 |
| *Eptesicus serotinus* | 24 | 18,60 | 33,00 | 24,8792 | 3,76078 |
| *Myotis myotis* | 236 | 19,90 | 39,90 | 28,2642 | 3,45991 |
| *Myotis nattereri* | 105 | 3,50 | 10,20 | 7,1829 | ,94822 |
| *Miniopterus schreibersii* | 1296 | 6,80 | 25,00 | 12,8472 | 1,83764 |

**Supplementary Table 3.**
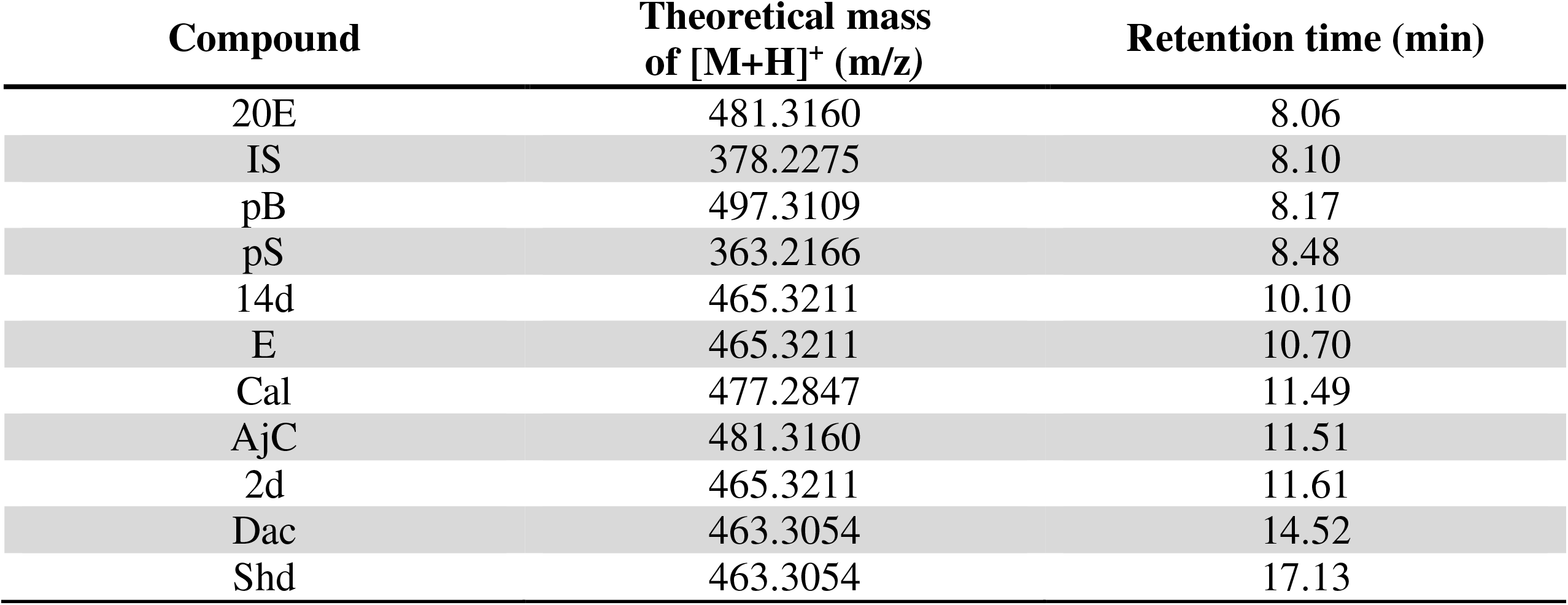
The main UHPLC-HRMS parameters of the targeted ecdysteroids and internal standard (IS).

## References

1. Munshi-South, J., and Wilkinson, G.S. (2010). Bats and birds: Exceptional longevity despite high metabolic rates. Ageing Res. Rev. 9, 12–19.

2. Hornok, S., Csorba, A., Kováts, D., Csörgő, T. and Hunyadi, A. (2019). Ecdysteroids are present in the blood of wild passerine birds. Sci. Rep. 9, 17002.

3. Hornok, S. et al.. (2016). An unexpected advantage of insectivorism: insect moulting hormones ingested by song birds affect their ticks. Sci. Rep. 6, 23390.

4. Dinan, L., and Lafont, R. (2006). Effects and applications of arthropod steroid hormones (ecdysteroids) in mammals. J. Endocrinol. 191, 1–8.

5. Parr, M.K. et al.. (2014). Estrogen receptor beta is involved in skeletal muscle hypertrophy induced by the phytoecdysteroid ecdysterone. Mol. Nutr. Food Res. 58, 1861–1872.

6. Feldhamer, G.A., Carter, T.C and Whitaker, Jr J.O. (2009). Prey consumed by eight species of insectivorous bats from southern Illinois. Am. Midl. Nat. 162, 43–51.

7. Catto, C.M.C., Hotson, A.M., Racey, P.A. and Stephenson, P.J. (1996). Foraging behaviour and habitat use of the serotine bat (Eptesicus serotinus) in southern England. J. Zool. 238, 623–633.

8. De Loof, A., Briers, T. and Huybrechts, R. (1984). Presence and function of ecdysteroids in adult insects. Comp. Biochem. Physiol. 79B, 505–509.

9. Rees, H.H. (2004). Hormonal control of tick development and reproduction. Parasitology 129 (Suppl.), S127–S143.

10. Luguterah, A. and Lawer, E.A. (2015). Effect of dietary guild (frugivory and insectivory) and other host characteristics on ectoparasites abundance (mite and nycteribiid) of chiropterans. Folia Parasitol. 62, 1–21.

## References

1. Estrada-Peña, A., Mihalca, A.D. and Petney, T.N. (Eds). (2017) Ticks of Europe and North Africa: A guide to species identification. Springer International Publishing, Cham, Switzerland, pp 404.

2. Kaňuch, P., Janečková, K. and Krištín, A. (2005). Winter diet of the noctule bat Nyctalus noctula. Folia Zool. 54, 53–60.

3. Bogdán, D. et al. (2018). Stereochemistry and complete 1H and 13C NMR signal assignment of C-20-oxime derivatives of posterone 2,3-acetonide in solution state. Magn. Reson. Chem. 56, 859–866.

4. Hunyadi, A. et al. (2007). Preparative-scale chromatography of ecdysteroids of Serratula wolffli andrae.J. Chromatogr. Sci. 45, 76–86.

5. Hunyadi, A. et al. (2016). Ecdysteroid-containing food supplements from Cyanotis arachnoidea on the European market: evidence for spinach product counterfeiting. Sci. Rep. 6, 37322.

6. Issaadi, H.M., Tsai, Y.C., Chang, F.R. and Hunyadi, A. (2017). Centrifugal partition chromatography in the isolation of minor ecdysteroids from Cyanotis arachnoidea. J. Chromatogr. B Analyt. Technol. Biomed. Life Sci. 1054, 44–49.

7. Agnarsson, I., Zambrana-Torrelio, C.M., Flores-Saldana, N.P. and May-Collado, L. J. (2011). A time-calibrated species level phylogeny of bats (Chiroptera, Mammalia). PLOS Currents Tree of Life. Edition 1.

8. Beck, A. (1995). Fecal analyses of European bat species. Myotis 32–33, 109-119.

9. Vaughan, N. (1997). The diets of British bats (Chiroptera). Mammal. Rev. 27, 77–94.

10. Presetnik, P. and Aulagnier, S. (2013). The diet of Schreiber’s bent-winged bat, Miniopterus schreibersii (Chiroptera: Miniopteridae), in northeastern Slovenia (Central Europe). Mammalia 77, 297–305.

11. Wilcoxon, F. (1992). Individual comparisons by ranking methods. In: Kotz, S., Johnson, N. L. (eds) Breakthroughs in statistics. Springer Series in Statistics (Perspectives in Statistics). Springer, New York, NY, 196–202.

12. Shapiro, S.S. and Wilk, M.B. (1965). An analysis of variance test for normality (complete samples). Biometrika 52, 591–611.

13. Benjamini, Y. and Hochberg, Y. (1995). Controlling the false discovery rate: a practical and powerful approach to multiple testing. J. Royal Stat. Soc. Ser. B 57, 289–300.

14. Gelman, A., Hill, J. and Vehtari, A. (2020). Regression and other stories. Cambridge University Press, Cambridge, UK, 548 pp.

15. R Core Team. (2021). R: A Language and Environment for Statistical Computing. Vienna, Austria: R Foundation for Statistical Computing. https://www.R-project.org/.

